# *In vivo* selection in non-human primates identifies superior AAV capsids for on-target CSF delivery to spinal cord

**DOI:** 10.1101/2023.09.13.557506

**Authors:** Killian S. Hanlon, Ming Cheng, Demitri De La Cruz, Nikita Patel, Miguel C. Santoscoy, Yi Gong, Carrie Ng, Diane M. Nguyen, Josette Nammour, Sean W. Clark, Karen Kozarsky, Casey A. Maguire

## Abstract

Systemic administration of adeno-associated virus (AAV) vectors for spinal cord gene therapy has challenges including toxicity at high doses and pre-existing immunity that reduces efficacy. Intrathecal delivery of AAV vectors into the cerebral spinal fluid (CSF) can avoid many of the issues of systemic delivery, although achieving broad distribution of the vector and transgene expression throughout the spinal cord is challenging and vector entry to the periphery occurs, sometimes initiating hepatotoxicity. Here we performed two rounds of *in vivo* biopanning in non-human primates (NHPs) with an AAV9 peptide display library injected intrathecally and performed insert sequencing on DNA isolated from either whole tissue (conventional selection), isolated nuclei, or nuclei from transgene-expressing cells. A subsequent barcoded pool of candidates and AAV9 was compared at the DNA (biodistribution) and RNA (expression) level in spinal cord and liver of intrathecally injected NHPs. Most of the candidates displayed enhanced biodistribution compared to AAV9 at all levels of spinal cord ranging from 2 to 265-fold. Nuclear isolation or expression-based selection yielded 4 of 7 candidate capsids with enhanced transgene expression in spinal cord (up to 2.4-fold), while no capsid obtained by conventional selection achieved that level. Furthermore, several capsids displayed lower biodistribution to the liver of up to 1,250-fold, compared to AAV9, providing a remarkable on target/off target biodistribution ratio. These capsids may have potential for gene therapy programs directed at the spinal cord and the selection method described here should be useful in clinically relevant large animal models.

## Introduction

Genetic diseases that affect the spinal cord are devastating, owing to their debilitating effects on quality of life as well as lack of effective treatments. Gene therapy for the central nervous system (CNS) using adeno-associated virus (AAV) vectors has risen to the forefront with promising clinical data showing efficacy of therapy targeting the CNS and spinal cord^1, 2^. Many of these approaches use systemic delivery of AAV serotypes such as AAV9 which can cross the blood-brain barrier. This generally gives reasonable distribution of transgene expression in neurons throughout the brain and spinal cord, although the efficiency with AAV9 and natural serotypes is fairly low and requires high doses of vector^3^. Systemic dosing also has major translational challenges such as pre-existing neutralizing antibodies to the AAV capsid, the high cost of generating the massive doses (>10^14^ vg/kg) required for therapy, and even dose limiting toxicities due to apparent complement activation and liver dysfunction^4, 5^. There is also concern of off target expression of transgene in dorsal root ganglion (DRG) neurons causing toxicity, as has been demonstrated in large animal models^6^, although this can be mitigated with miRNA transgene expression detargeting strategies^7^.

Therefore, research has been focused on developing AAV-based gene therapies using direct injection of vector into the cerebral spinal fluid (CSF). Intrathecal injection into the cerebral spinal fluid (CSF) around the lumbar spinal cord with AAV generally gives good transduction of motor neurons in the lumbar region, but less transduction of cervical regions and the brain^8, 9^. Positioning the subject in the Trendelenburg position can enhance vector biodistribution of the cervical region somewhat^9, 10^. On the other hand, cisterna magna injection of AAV vectors gives better cervical and brain transduction but less so in the lumbar region^11^. There is also a surprisingly high amount of vector that enters the circulation after CSF delivery which ends up in the liver^12, 13^, which could lower the safety benefits of a more local delivery. In fact, in a recent study by Hudry et al., the team demonstrated liver toxicity in cynomolgus monkeys after intrathecal delivery^14^. And in a clinical trial for spinal muscular atropy (SMA), 2 out of 25 patients receiving intrathecally administered AAV9-sc-CBA-SMN1 (onasemnogene abeparvovec; Zolgensma) had liver enzyme elevations^15^. There is a clear need for AAV vectors that can transduce the spinal cord with the following properties: (1) high efficiency at lower dose, (2) distribution throughout the entire spinal cord from lumbar to cervical regions, (3) low off target leakage/transduction of peripheral organs, especially liver.

With this in mind, we developed an *in vivo* selection strategy with an AAV9 peptide display library (iTransduce) that allows selection of transduction competent AAV capsids^16^. We further developed the system to be utilized in non-human primates (NHPs) to maximize clinical translation of the capsids’ transduction properties, owing to physiological and anatomical (e.g. size, CSF volume) differences between primates and mice. We performed two rounds of *in vivo* selection in NHPs after intrathecal injection of the AAV peptide display library. In the second round, we utilized iTransduce to isolate AAV variants that led to functional transduction. We then identified capsids which outperformed AAV9 in terms of biodistribution and transduction efficiency in the spinal cord and lowered biodistribution to liver.

## Results

### *In vivo* intrathecal injection AAV library selection strategy in NHPs

We have previously reported on the use of the iTransduce AAV peptide display library. The genome construct consists of a promoter driving a Cre-recombinase cassette as well as p41 promoter-driven AAV9 capsid gene with 7-mer peptide inserts between amino acids 588-589 of VP1. This allows surface display of 50 copies of peptides on VP3 on the capsid surface. For round one of our *in vivo* selection strategy in NHPs, we produced the AAV peptide display library and injected it intrathecally into Old World monkeys (cynomolgus macaques) (**Fig. 1a**, **Table I**). Next, AAV genome DNA was recovered by PCR from cervical, thoracic, and lumbar regions of the spinal cord and subjected to next generation sequencing (NGS) to analyze the content of the 7-mer encoding inserts. These capsid inserts were pooled and packaged into the round 1 selected spinal cord library (**Fig. 1a**). For the second selection round, we developed a strategy to enable selection of capsids that could mediate transduction expression in a non-transgenic animal such as an NHP. To do this we used a two-vector approach as outlined in **Fig. 1b**. The first vector is the AAV iTransduce library genome shown in **Fig. 1a**, that is packaged inside the round 1 rescued capsid inserts. The second vector is a Floxed reporter cassette encoding an H2B-fused mPlum protein called AAV-CBA-Floxed-STOP-H2B-mPlum. We used H2B-mPlum (mPlum with a nuclear localization signal) to allow sorting of fluorescent nuclei from either fresh or frozen tissue (cytoplasmic proteins are likely to leak out of freeze/thawed tissues). When the AAV capsid library encoding Cre is co-injected or sequentially injected with the AAV9-CBA-floxed-STOPH2B-mPlum, cells that are co-transduced with library capsids able to express Cre along with the AAV9 capsid will excise the stop cassette and express H2B-mPlum in the nucleus (**Fig. 1b**). Spinal cord is isolated and DNA can be purified from whole frozen tissue, purified nuclei, or H2BmPlum flow-sorted nuclei. Peptide inserts are analyzed by NGS and candidate AAV capsids chosen for further characterization.

**Figure 1.**
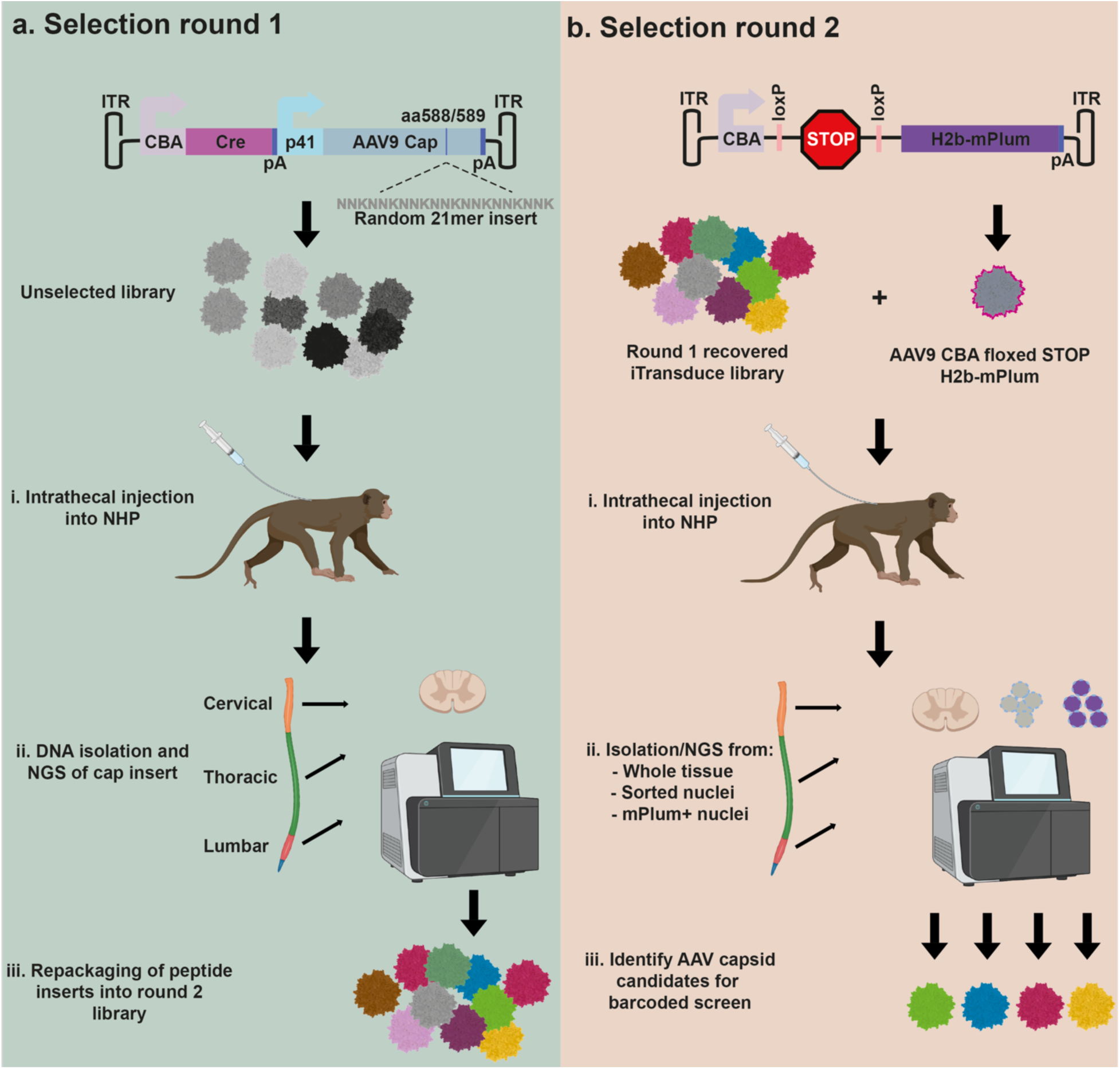
Overview of the *in vivo* selection in non-human primates (NHPs) to identify AAV capsids with the ability to efficiently transduce spinal cord. **a.** Round 1 of selection. The iTransduce AAV genome plasmid has a CBA-driven Cre cassette followed by a p41 driven AAV9 capsid with randomized 21mer bp inserts encoding for 7mer peptides inserted into the capsid. NHPs are injected intrathecally (IT) with the library and three weeks later animals are sacrificed and spinal cord removed. Next generation sequencing (NGS) is performed on AAV genomes containing 21mer inserts amplified from three spinal cord regions. The rescued cap inserts are inserted into the iTransduce backbone and packaged into the R1 rescued library. **b.** Round 2 of selection. A second reporter AAV expression plasmid is used to allow transduction-based selection in a non-transgenic animal. The reporter contains a Cre-sensitive floxed-STOP-H2B-mPlum cassette. When co-injected intrathecally into NHPs with iTransduce library from Round 1, cells that are co-transduced with Cre-expressing capsids can rescue nuclear mPlum expression. Spinal cord is again isolated and DNA isolated using three different strategies: (1) whole tissue DNA as in 1a, (2) flow-sorted nuclei, (3) flow-sorted H2B-mPlum nuclei (contains transduction-competent capsid DNA). NGS is performed and AAV capsids are identified for further screening for spinal cord transduction.

**Table I.** NHP study information.

| Animal ID | Age (years, months) | Weight (kg) | Sex | Vector | Dose (VG) | AAV NAb titer |
| --- | --- | --- | --- | --- | --- | --- |
| M682 | 1 yr, 8 mo. | 2.1 | Male | Unselected library (Round 1) | $9.0 \times 10^{11}$ | <5 |
| N495 | 1 yr, 11 mo. | 1.9 | Female | Round 1 recovered library (Round 2) | $2.0 \times 10^{11}$ | 5 |
| MB1255 | 5 yrs, 5 mo. | 5.4 | Male | Barcoded library candidates (screen) | $4.48 \times 10^{12}$ | <5 |
| EC1042 | 5 yrs, 2 mo. | 5.6 | Male | Barcoded library candidates (screen) | $4.48 \times 10^{12}$ | <5 |

### Round 1 of selection of intrathecally injected AAV9 peptide display library in NHPs

A preparation of the iTransduce AAV9 peptide display library was produced, purified, and titered. Library diversity was determined to be adequate for the selection and a male cynomolgus monkey was injected intrathecally with 9×10^11^ vector genomes (vg) of the AAV library. Three weeks later, the animal was killed, perfused with sterile saline and tissues including the spinal cord were flash frozen and stored at - 80°C. We isolated DNA from homogenized, whole, spinal cord samples from the cervical, thoracic, and lumbar regions. Next, we performed a PCR with primers surrounding the 21-mer inserts encoding the 7-mer peptides and submitted the three samples for low-depth NGS. We obtained diverse inserts from the sequencing data; reads and unique inserts frequencies are shown in **Table II**.

**Table II.** Round 1 sequencing reads of AAV capsid inserts in spinal cord DNA.

| Spinal Cord Region | Unique variants (insert reads) | Unique sequences (individual 21-mer inserts) |
| --- | --- | --- |
| Lumbar | 57,000 | 24,000 |
| Thoracic | 62,000 | 21,000 |
| Cervical | 25,000 | 1,600 |

Read diversity (number of unique reads) in the cervical sample was much lower than thoracic or lumbar. This is likely due to few capsids with the ability to traffic from the lumbar region to the cervical region after intrathecal injection. We pooled amplified capsid DNA fragments containing 21-mer inserts from all three spinal cord regions and ligated them into the iTransduce plasmid library backbone.

### Two vector system allows detection of transduction competent AAV capsids in non-transgenic large animals

To create a system that would allow us to detect transgene expression at the protein level of transduction-competent AAV library variants, we designed a second vector, AAV-CBA-Floxed-STOP-H2B-mPlum as described in **Fig. 1b**. To test the function of the system we transduced cultured 293T cells with AAV capsids packaging the following expression cassettes: (1) AAV-PHP.B-CBA-Cre only, (2) AAV9-floxed-STOP-H2B-mPlum only, (3) a mixture of AAV-PHP.B-CBA-Cre with AAV9-floxed-STOP-H2B-mPlum. Three days later, cells were examined by fluorescence microscopy for both DAPI and H2B-mPlum fluorescence. As expected, nuclear-localized mPlum fluorescence was detected only in the cells co-transduced with both vectors (**Fig. 2a**). To test whether the system worked in non-transgenic animals *in vivo*, we intravenously injected wild type C57BL/6 mice with the following groups of vectors: (1) AAV-F-CBA-Cre only, (2) AAV-F-Floxed-STOP-H2B-mPlum only, or (3) a mixture of AAV-F-CBA-Cre with AAV-F-Floxed-STOP-H2B-mPlum. We used the previously described AAV-F capsid as it efficiently transduces the brain after systemic injection^16, 17^. Sixteen days post injection, mice were perfused with PBS and brains flash frozen. Next, we isolated nuclei from dissociated brain, labeled total nuclei with a violet-fluorescing dye, and then analyzed nuclei from each group from mPlum fluorescence (**Fig 2b**). All groups had nuclei, as observed by the violet dye stain. However, only the AAV-Floxed-STOP-H2B-mPlum+ AAV-Cre group showed many fluorescent mPlum+ nuclei, which demonstrates the specificity and functionality of the two-vector system (**Fig. 2c**).

**Figure 2.**
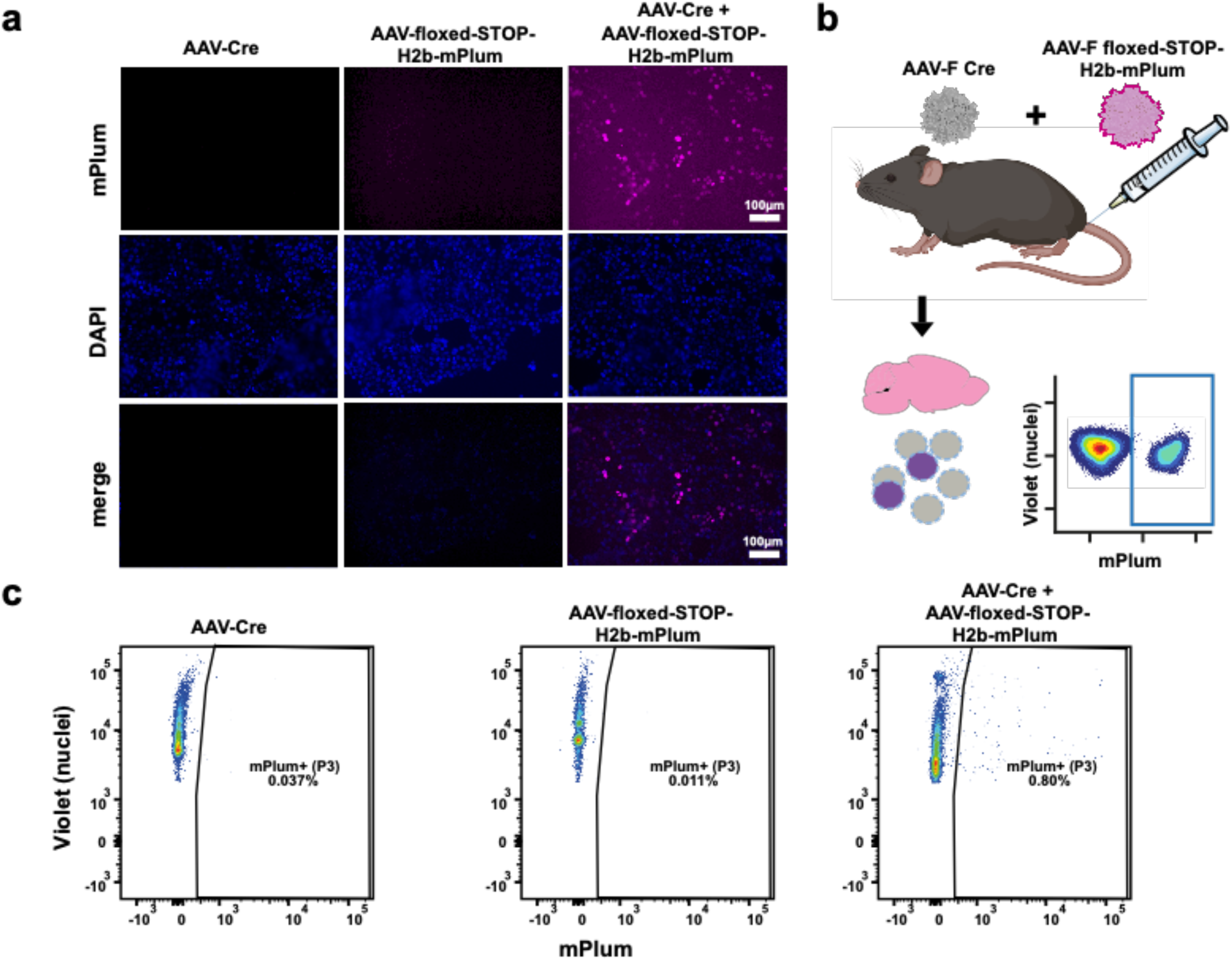
Two-vector iTransduce system allows detection of transduction competent AAV *in vitro* and *in vivo.* **a.** 293T cells were transduced with AAV-Cre or AAV-Floxed-STOP-H2B-mPlum vectors alone (first two columns) or co-transduced with both vectors (far right column). Nuclear H2B-mPlum fluorescence was only detected in co-transduced cells. Scale bar= 100 µm. **b.** Wild type C57Bl/6 mice were iv injected with the following vector groups (n=2 mice/group): (1) AAV-Floxed-STOP-H2B-mPlum, (2) AAV-Cre, or (3) a mixture of both vectors in 1 and 2. **c.** Brains were harvested, frozen, and nuclei isolated and then labeled with violet dye and analyzed with flow cytometry for violet and H2B-mPlum.

### Round 2 of selection enriches for capsids that are maintained in the NHP spinal cord after intrathecal injection

For the second round of selection, we performed two separate intrathecal injections into a single female cynomolgus macaque. The animal was first intrathecally injected with 2×10^11^ vg of the recovered iTransduce library from round one and three hours later, while the animal was still anesthetized, intrathecally injected with 3×10^13^ vg of AAV9-CBA-Floxed-STOP-H2B-mPlum. Three weeks later the animal was killed, and spinal cord and other tissues collected and flash frozen. As shown in **Fig. 1b** we isolated AAV genomes containing the 21 bp inserts via three methods: (1) whole tissue as in round 1, (2) sorting dye-labeled nuclei from dissociated spinal cord, and (3) sorting H2B-mPlum-positive nuclei from cells co-transduced by a transduction competent AAV capsid (Cre-expressing) and the AAV9-Floxed-STOP-H2B-mPlum vector. For whole tissue isolated DNA, we used samples from all three regions of the spinal cord as in round 1. For the nuclei isolation, we used the lumbar region of the spinal cord to increase the chances of detecting mPlum expression as this was the injection region of vector and should have the highest vector concentration. We purified nuclei using iodixanol density gradients, labeled the nuclei using violet dye and flow sorted the nuclei in both mPlum negative and mPlum-positive fractions. We set the mPlum+ nuclei gate based on a round 1 spinal cord sample which was not injected with the AAV9-Floxed-STOP-H2B-mPlum vector (**Fig. 3a**). mPlum+ nuclei were detected in the co-injected animal **(Fig. 3b**), and we obtained 2.09×10^6^ mPlum-nuclei and 1.70×10^4^ mPlum+ nuclei. For the whole tissue isolated DNA, and the nuclei from both mPlum+ and mPlum-fractions, we were able to amplify the insert-containing PCR product. All PCR amplicons were analyzed by NGS. For the whole tissue DNA samples isolated from thoracic and lumbar spinal cord regions we observed many of the same top peptides from round 1. There was strong enrichment however, with top peptides showing up to 15-fold to 21-fold enrichment compared to the round 1 recovered library (**Supplementary Table**). The read frequency of the top clones was 0.25% and 0.28% for lumbar and thoracic, respectively. For the cervical sample there was an even greater enrichment in round 2 recovered sequences compared to the round 1 recovered library. The top peptides were enriched by 25 to 67-fold (**Supplementary Table**). For the mPlum-positive and negative nuclei there were far fewer variants. The top peptide in the mPlum+ fraction, HTPLPRP, represented 88% of reads and the mPlum negative was 17% of reads. Due to the surprising level of read frequency of one peptide in the mPlum-positive nuclei fractions, we repeated the PCR on the nuclei DNA template for a total of three times. Interestingly in the two repeat runs of the mPlum+ fraction, the top peptides, PKYPLLG and RPDHVRK, were both at high frequencies (>99% of reads). Based on these findings of different peptides dominating the NGS reads in the same spinal cord sample but in independent PCR reactions, we assumed that we were observing stochastic effects of the PCR owing to the low amount of nuclei present for template in the PCR. However, we considered all of these top peptides as viable candidates as they were found in the mPlum+ fraction.

**Figure 3.**
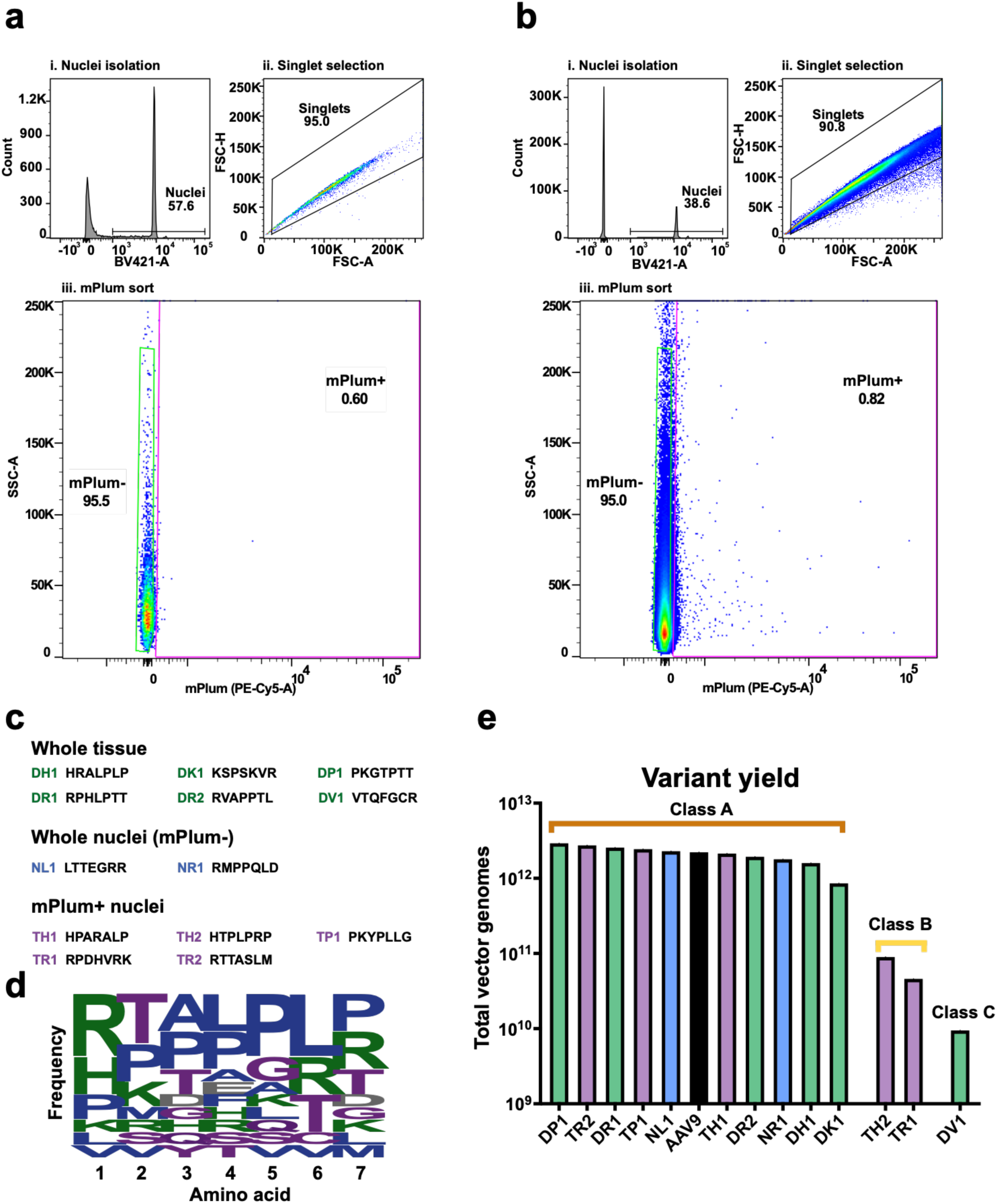
Isolation and selection of candidate variants in NHPs. **a-b.** Flow cytometry of nuclei isolated from control (a) and library-injected (b) NHPs. Nuclei were selected using fluorescent dye, with mPlum used to discriminate nuclei from transduced cells. **c.** Candidate variants were selected from whole tissue isolation, mPlum-nuclei, and mPlum+ (transduced) nuclei. The names and insert peptides of each variant are listed. **d.** The frequency of amino acids at each position for the selected candidates is given. Larger size indicates more frequent incorporation of a given amino acid, for a given position. **e.** Overall production yield of each capsid variant when individually produced, compared to AAV9 (in black).

To choose candidates for further screening, we included the following parameters: (1) For the whole tissue isolated DNA data, we chose peptides that showed up in all three regions of spinal cord to identify capsids that mediate good biodistribution. (2) We included peptides that were enriched in the cervical region of the spinal cord to increase chances of having capsids that can traffic from lumbar to cervical region. (3) We included candidates from the nuclei sorted samples (mPlum-positive and negative) to increase chances of having transduction competent capsids. (4) We looked for common motifs in the peptides. Based on these parameters we chose 13 candidate peptides for further analysis (**Fig. 3c and Table III**). An amino acid frequency calculator showed a characteristic LPLP motif at positions 4-7 (**Fig. 3d**). We next tested the yield of individually produced candidates (in addition to AAV9 for comparison) to understand if they would be manufacturable for further development. The majority of capsids (10/13) produced very well, better than or very close to the yield of AAV9 (**Fig. 3e**). Three other capsids produce at lower levels. We classed these capsids A, B, and C, based on their production efficiency, with A having the highest and C the lowest (**Fig. 3e**).

**Table III.** Barcoded spinal cord candidate capsids for screen.

| Candidate Name | Peptide Sequence | Selection Type | Reason for choosing |
| --- | --- | --- | --- |
| **DK1** | **KSPSKVR** | whole tissue | Top peptide in Cervical region |
| **DP1** | **PKGTPTT** | whole tissue | PKX motif, high frequencing in lumbar and thoracic regions |
| **DR2** | **RVAPPTL** | whole tissue | Highest enrichment in thoracic region, high in lumbar region |
| **DR1** | **RPHLPTT** | whole tissue | LP motif at 4,5 position, RP motif, enriched in cervical region |
| **DH1** | **HRALPLP** | whole tissue | LP motif at 4,5 position |
| **DV1** | **VTQFGCR** | whole tissue | Top peptide in Cervical region |
| **NL1** | **LTTEGRR** | nuclei | High frequency |
| **NR1** | **RMPPQLD** | nuclei | High frequency |
| **TR2** | **RTTASLM** | mPlum+ nuclei | Detected at low frequency in mPlum+ reads |
| **TH1** | **HPARALP** | mPlum+ nuclei | Detected at low frequency in mPlum+ reads; similar sequence to whole tissue peptide HRALPLP |
| **TP1** | **PKYPLLG** | mPlum+ nuclei | PKX motif, Top frequency read |
| **TR1** | **RPDHVRK** | mPlum+ nuclei | Common RP motif, Top frequency read |
| **TH2** | **HTPLPRP** | mPlum+ nuclei | LP motif at 4,5; RP motif, Top frequency read |

### Barcoded capsid screen reveals variants with enhanced biodistribution in spinal cord and lowered liver biodistribution compared to AAV9

To assess the potential of the candidate peptides identified from round 2 of selection to mediate transduction of spinal cord, we engineered an AAV transgene expression cassette shown in **Fig. 4a**. We used a human frataxin cDNA fused to a hemagglutinin tag similar to Goertsen et al^18^, as this construct has shown to express well in NHP CNS and potentially avoid confounding effects of immunogenic/toxic fluorescent proteins like GFP. We added a barcode region which allows us to differentiate each of our candidate capsids via NGS at both the DNA and RNA levels. We tested the barcode system to measure biodistribution and transduction differences between AAV capsids by comparing NGS reads from barcodes at the DNA and RNA levels using systemically injected pooled AAV9 and AAV-F^16^ vectors (each with their own barcode) in C57BL/6 mice. Based on our prior work, the AAV-F capsid has been shown to mediate enhanced biodistribution and transduction of brain compared to AAV9, so we expected the barcode data to yield a similar profile if it was performing as expected. Similar to our prior data with single capsid comparisons^16^, AAV-F mediated a 13-fold increase and 28-fold increase in AAV genomes and encoded RNA, respectively (**Supplementary Fig. S1**).

**Figure 4.**
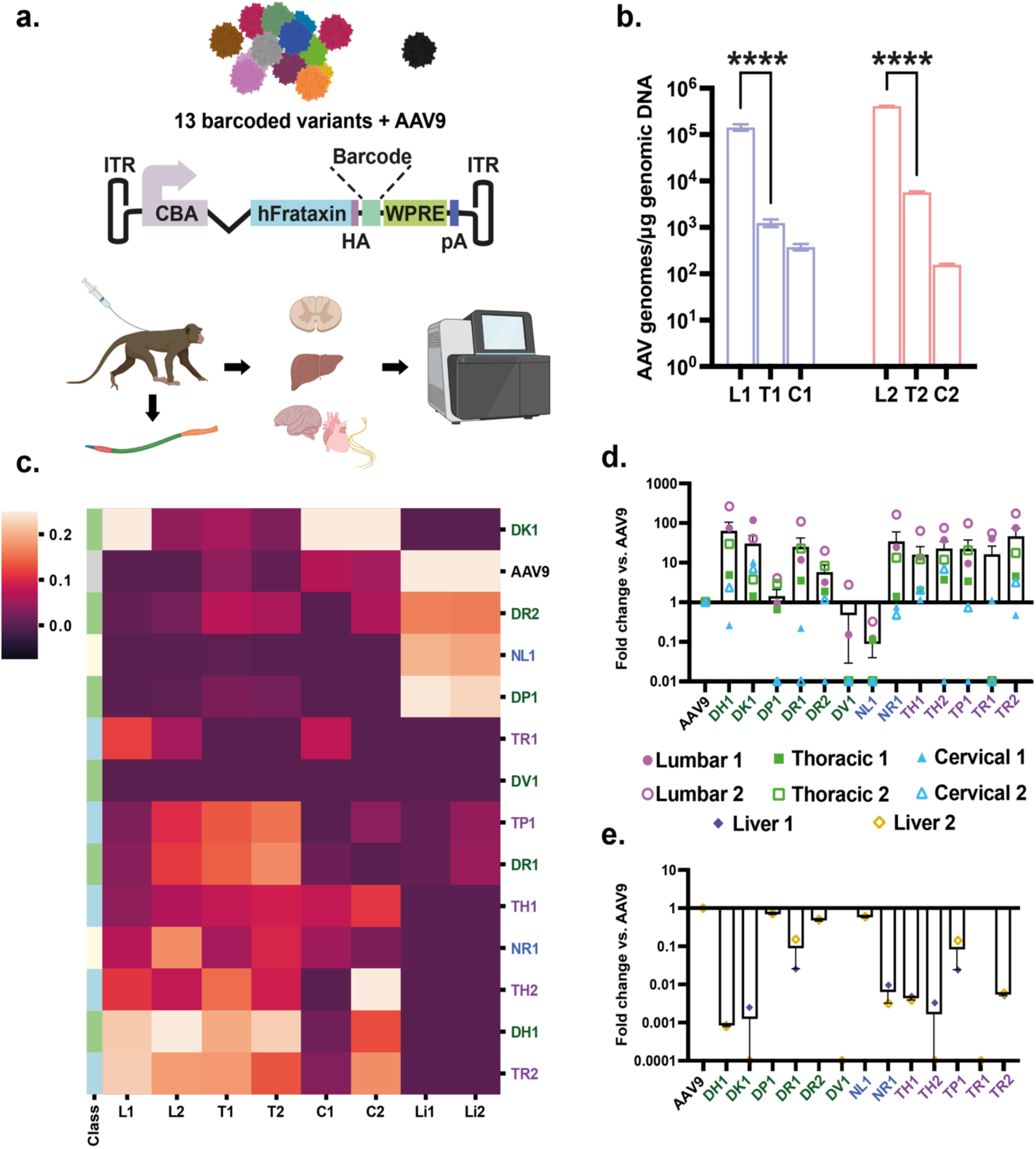
Barcoded candidate capsid screen identifies variants with enhanced biodistribution in NHP spinal cord and reduced biodistribution to liver. **a.** Schematic showing the procedure used for this experiment. Each capsid variant was barcoded between the protein-coding region and post-transcriptional sequences. Following intrathecal injection, DNA wasisolated from 1) each region of the spinal cord, 2) liver, 3) brain, 4) peripheral nerves, and 5) heart. These were subjected to next-generation sequencing. **b.** Overall biodistribution of AAV capsid genomes in each section of the spinal cord as measured by qPCR. L1, T1, C1 are from NHP #1001. L2, T2, C2 are from NHP #1002. **C.** Heatmap showing relative frequency of each variant (or AAV9) in each region of the spinal cord and liver for each animal. Variants are clustered by expression pattern (higher expression= lighter color). **d-e.** Average spinal cord (d) and liver (e) biodistribution of each variant expressed as fold-change vs. AAV9. Zero values are not displayed owing to logarithmic scaling of the graphs. Error bars represent standard error. L, T, C (L1= NHP #1001, L2= NHP #1002, etc.): Lumbar, Thoracic, Cervical regions of the spinal cord. Li (Li1, 2) = Liver samples for #1001 and #1002, respectively. Color coding for the different capsids: Green= whole tissue isolated; Blue= nuclei isolate; Purple= mPlum-positive nuclei isolated.

For the experiment in NHPs, we individually produced each of the 13 capsid candidates as well as AAV9, which served as our benchmarking capsid (**Fig. 4a**). Next, we pooled the library candidates as well as AAV9. Class A capsids (which includes AAV9) were at a dose of 4×10^11^ vg/capsid, Class B at a dose of 2×10^10^ vg/capsid, and Class C at 3.98×10^9^ vg. Two adult male cynomolgus monkeys (**Table I**) were each injected IT with 4.48×10^12^ vg of the pooled barcoded capsids. Three weeks later, animals were killed, and spinal cord and other tissues isolated. We isolated DNA and RNA from the cervical, thoracic, and lumbar regions of the spinal cord and DNA and RNA from the liver. To understand the level of the pooled capsid genome in the spinal cord, we performed qPCR which detects AAV genomes using primers and probes to the poly A signal of the expression cassette. For both NHPs, AAV genomes were highest in the lumbar region with a steep gradient from lumbar to thoracic and further drop-off to cervical spinal cord (**Fig. 4b**), which is expected given the low dose of administered vectors and that it was a pool of different capsids. To understand the frequency of each capsid at each level of the spinal cord as well as from the liver, NGS was performed using the barcode unique for each capsid. As can be observed in the heat map of the NGS data, many capsids displayed higher frequency than AAV9 in all three spinal cord regions, with some variability between animals (**Fig. 4c**). Remarkably, 10 capsids outperformed AAV9 in levels of vector genomes in the lumbar region, ranging from 2 to 265-fold (**Fig. 4d**). Several capsids had higher levels of genomes in thoracic and cervical regions compared to AAV9, although there was more inter-animal variability compared to the lumbar region (**Fig. 4d**). Notably, biodistribution to the liver was reduced for the majority of tested capsids (**Fig. 4c**), with ranges of 2- to 1,250-fold lower than AAV9 (**Fig 4e**). We also performed barcode analysis on the capsids in four brain regions, dorsal root ganglia (three levels of spinal cord), and heart for both NHPs (**Supplementary Fig. S2**). There was a high degree of variability between capsids and inter-animal differences for the same capsid in some cases. Some capsids, such as DH1, showed a much higher read frequency in most brain samples compared to AAV9. Finally, we performed barcoded frequency analysis for each of the capsids in the following nerves isolated from the two NHPs: sciatic nerve, 8^th^ cranial nerve, sural nerve, and ulnar nerve (**Supplementary Fig. S3**). While there was variability between the two animals for some capsids, there were some trends for increased frequency in certain nerves over AAV9. For examples capsids DP1 and DR2 had high read frequencies in sural and ulnar nerves compared to AAV9, DH1 had higher levels in sciatic nerve and 8^th^ cranial nerve, and DK1 in 8^th^ cranial nerve.

### Enhanced transduction in spinal cord and lower transduction of liver with several candidate capsids

To assess expression levels of the capsids, we assessed levels of cDNA reverse transcribed from frataxin-HA mRNA using RT-qPCR. The cDNA levels were detected in both animals in the lumbar region, although the levels were significantly higher (12.5-fold) in NHP #1001 than #1002 (**Fig. 5a**). This could be due to differences in transduction efficiency between animals, as the vector genomes were quite similar between animals (**Fig 4b**). The levels of cDNA in thoracic and cervical regions were not above the negative controls, likely owing to the low injected dose of each capsid, so further analysis of mRNA/cDNA was focused on the lumbar region of spinal cord and the liver. NGS was performed on the barcode cDNA from mRNA isolated from lumbar spinal cord and liver. We obtained high numbers of reads from lumbar region of both animals and frequencies of top capsids were relatively consistent between independent PCR runs (four runs/animal) (**Fig. 5b**). In contrast to the DNA biodistribution data, at the RNA level fewer capsids outperformed AAV9. None of capsids from the whole tissue DNA isolation outperformed AAV9 in at least one of the injected animals by a factor of ≥2. In contrast, 1 of 2 in the nuclei isolation did, and 3 of 5 in the mPlum+ nuclei isolation did (**Fig. 5b**). The average enhancement ranged from 1.8 to 2.4-fold for the four capsids over AAV9 and up to 3.5-fold in one of the two animals (**Fig. 5b**). Liver RNA expression appeared similar to the DNA biodistribution data; capsid variants showed expression levels 2 to 30,000-fold lower compared to AAV9 (**Fig. 5c**).

**Figure 5.**
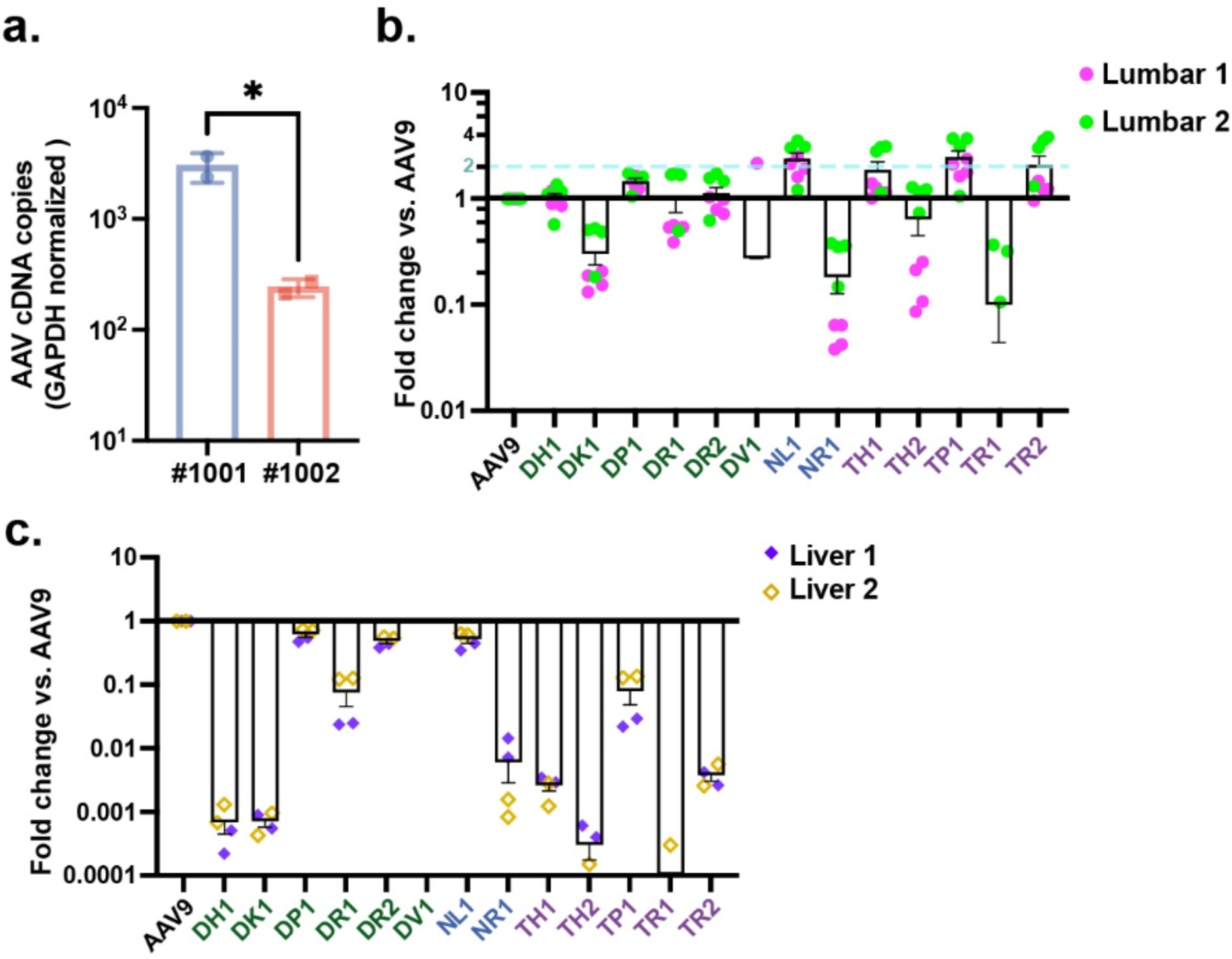
Barcoded capsid screen identifies variants with enhanced transgene RNA levels in spinal cord and reduced RNA levels in liver. **a.** Expression levels of the AAV transgene in the lumbar portion of the spinal cord of each NHP (#1001, #1002), relative to GAPDH mRNA. **b-c.** Average RNA expression values for the lumbar region of the spinal cord (b) and liver (c), expressed as fold-change vs. AAV9. Zero values are not displayed owing to logarithmic scaling of the graphs. Error bars denote standard error. Lumbar 1= NHP #1001; Lumbar 2= NHP #1002. Liver 1= NHP #1001; Liver 2= NHP 1002.

### Generating a profile of candidate capsid features allows identification of top candidates for preclinical development

Based on the data in NHPs from biodistribution in spinal cord and liver and transduction in spinal cord as well as production efficiency, we selected the top candidate capsids to move forward with (**Table IV**). This allowed us to narrow 13 initial candidates down to 4 peptides, which were peptides NL1, TH1, TP1, and TR2. It is important to note that all chosen capsids were from the higher-yield class A capsids injected at the same dose as AAV9.

**Table IV.** Top spinal cord capsid candidate features.

| Variant name | Insert | Spinal cord biodistribution (L-T-C) | Spinal cord transduction (Lumbar) | Liver biodistribution and transduction | Production efficiency |
| --- | --- | --- | --- | --- | --- |
| NL1 | LTTEGRR | Low | Moderate | Low | High |
| TH1 | HPARALP | High | Moderate | Low | High |
| TP1 | PKYPLLG | High | Moderate | Low | High |
| TR2 | RTTASLM | High | Moderate | Low | High |

### NHP-selected capsids efficiently transduce murine spinal cord after intrathecal injection

Since the selection of the library was performed in NHPs, it’s likely the transduction profile would be more efficient in this species compared to rodents. However, since most neurological models of human disease are developed in transgenic mice, it would be convenient for preclinical development if the NHP-selected capsids were also functional in mice. To assess whether the AAV capsids selected in NHP were “backwards compatible”, adult 6-8 week old C57BL/6 mice were injected intrathecally in the lumbar region individually with either AAV9, NL1, TH1, TP1, or TR2 (n=5/capsid). AAV capsids packaged a single-stranded AAV-CBA-eGFP genome. AAV9, TH1, TP1, and TR2 were dosed at 8.3×10^10^ vg/mouse and NL1 at 4.8×10^10^ vg/mouse, owing to a lower titer obtained for this capsid. Four weeks post injection, mice were euthanized, and spinal cords and livers were harvested and cryosectioned. Spinal cords were divided into rostral and caudal regions and immunostained for GFP followed by immunohistochemical detection. Cell types were identified by morphology and scored separately for neurons, glia, pia mater cells, and ependymal cells surrounding the central canal. All capsids transduced the murine spinal cord after intrathecal injection (**Fig. 6a**). A semi-quantitative data analysis was performed whereby transduction percentages were divided amongst five grading categories, with one being the lowest and five the highest percentage of GFP-positive cells (**Fig. 6b**). Scoring for transduction percentages in both rostral and caudal spinal cord revealed that all capsids were able to transduce both neurons and glia, with some intragroup variability in grading (**Fig. 6b, c**). Despite being injected at 58% of the dose of the other capsids, NL1 generally trended towards higher percentages of transduced neurons in more mice/group compared to the other capsids and AAV9 (**Fig. 6b,c)**. Glial cells were also readily transduced (**Fig. 6b, d**). All capsids were also able to transduce the pia mater and ependymal cells (data not shown). Livers were also examined for GFP expression based on our data in NHPs that showed lower biodistribution to this organ after intrathecal injection of the pooled candidates. Interestingly, in contrast to the NHP data, in mice the novel capsids transduced similar percentages of liver cells compared to AAV9 (**Supplementary Fig. S4**).

**Fig 6.**
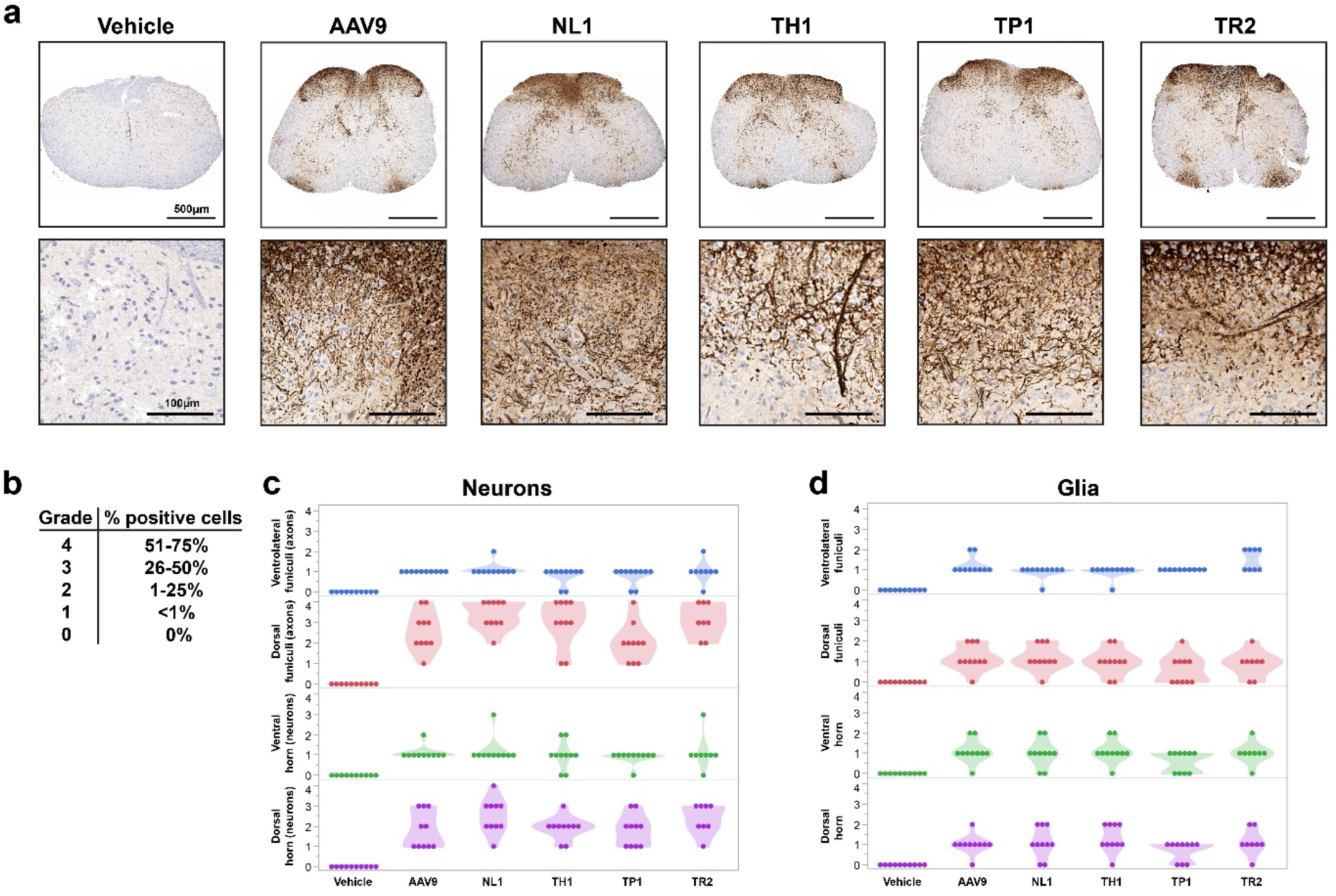
NHP selected AAV capsid candidates transduce neurons and glia in mice after lumbar intrathecal bolus injection. Mice (n=5/capsid) were injected intrathecally with AAV9, NL1, TH1, TP1, and TR2 packaging an AAV-CBA-GFP transgene cassette. All capsids were injected at 8.3×1010 vg/mouse with the exception of NL1, which was dosed at 4.8 x1010 vg/mouse **a**. Representative low and high magnifcation images of spinal cord for each capsid. Brown DAB (3,3’-Diaminobenzidine) staining indicated GFP immunoreactivity. **b.** Score grading for spinal cord transduction efficiency. **c.** Transduction scoring for neurons in spinal cord. **d.** Transduction scoring for glial cells in spinal cord. Each data point represents either a lumbar or cervical region for each mouse analyzed.

## Discussion

While AAV9 has been successfully developed for transgene delivery to the spinal cord to treat the neuromuscular disease, spinal muscular atrophy (SMA), there is room for improvement. When delivered at high systemic doses, AAV vectors can cause liver enzyme elevation, complement activation, thrombotic microangiopathy (TMA), and even sepsis, organ failure, and death^4, 19^. Thus, much recent research is focused on developing AAV gene therapies for CNS disorders using a CSF route of administration.

Delivery of AAV directly into the CSF has been shown to reduce exposure to neutralizing antibodies which are present at much lower levels than in blood^20^. While there is less systemic exposure to vector than a systemic route of administration, there is still a large amount of vector that can enter the blood and peripheral organs after CSF injection. A recent study by Meseck et al. performed lumbar IT injection of NHPs and measured vector genomes in CNS and peripheral tissues 4 weeks post injection^12^. Remarkably, liver received over 100 times the number of AAV genomes compared to the brain and approximately 10 times more than the spinal cord after lumbar intrathecal injection^12^. Furthermore, transgene expression was higher in the liver and heart compared to spinal cord^12^. Other studies using radiolabeled AAV capsids injected into the CSF of NHPs – including AAV9 – has demonstrated that up to approximately 90% of administered capsid can enter the periphery^13, 21^. Obviously, this high off-target biodistribution runs completely counter to the intent of a direct CSF administration. Transduction of liver after intrathecally injected AAV is more than just a theoretical risk for toxicity. Recently Hudry et al. found that both intravenously and intrathecally injected AAV9 encoding a self-complementary GFP or SMN1 transgene cassette resulted in hepatotoxicity as measure by liver enzyme elevations and chemistry, histopathology, cytokines, and complement activation^14^. Immunosuppression with Prednisolone_ or Rituximab/Everolimus were not able to mitigate these effects, suggesting that current pharmaceuticals will not suffice^14^. Interestingly, the researchers found that AAV transduction needed to occur, as empty capsids and promoterless transgene cassettes did not result in toxicity^14^. The most straightforward way to avoid transgene expression in the liver is to either avoid biodistribution to liver using engineered capsids like the ones described in our study, or restrict transgene expression away from the liver using tissue specific promoters^22^ or miRNA based transgene mRNA degradation^23, 24^.

Our work is, to our knowledge, the first selection of an AAV peptide display library in NHPs using the intrathecal route. Remarkably, the majority of the top capsids identified in our selection showed greatly reduced biodistribution (up to 1,250-fold less frequent reads) and transduction (up to 30,000-fold less frequent reads) in liver compared to AAV9. This desirable feature may allow for less toxicity observed with liver transduction, although this will need to be confirmed by testing with individual capsids encoding transgenes of interest in toxicology studies in NHPs. Furthermore, the top candidate capsids appear to be more efficient than AAV9 at spinal cord transduction at the RNA level (**Fig. 5b**). This latter outcome may have been influenced by the selection approach. To enable detection of transgene expressing cells during the selection rounds, we made some modifications to our prior reported iTransduce system^16^ to allow its use in non-transgenic animals. To do this we used a two-vector system with the library encoding Cre recombinase as usual and the second vector packaging the Cre inducible reporter (Floxed-STOP-H2B-mPlum). Using this system, we were able to flow sort H2B-mPlum fluorescent nuclei that applied additional selective pressure to identify transduction-competent capsids. The reporter capsid (AAV9 in our case) likely influences the selection outcome, as fluorescent nuclei are restricted to cells which can be transduced by AAV9. In our study, this was a benefit, as we wanted to maintain the tropism of AAV9 in the spinal cord (glia and neurons), while improving transduction efficiency and biodistribution. However, if other cell types outside the tropism of AAV9 are desired as targets, this approach may not be feasible, or a different capsid packaging the reporter gene which can transduce the target cell would need to be utilized.

We made several interesting observations during the study. First, many of the peptides had a leucine:proline (LP) motif in the 7-mer sequence, which may be important in their enhanced biodistribution in the spinal cord compared to AAV9 (**Fig. 4d**). Second, we found that the capsids isolated from nuclei or from nuclei in transduced cells had higher levels of transduction (RNA-based reads) compared to capsids identified from the whole-tissue isolated DNA (**Fig. 5b**). Whole tissue isolation is likely to contain AAV genomes that may be outside the cell (still in capsids) or nucleus or not uncoated in the nucleus (i.e., not transcriptionally active). On the other hand, nuclear isolation and transgene-based selection is at or close to the final steps of transduction. While this observation was based on a limited number of capsids from each method, owing to a limit of capsids that we could test by barcoding, it does suggest, in line with other reports^16, 25-27^, that selection as far down stream in the transduction process as technically feasible is likely to yield capsids capable of transduction.

While we did observe capsids that appear more efficient than AAV9 at both the biodistribution (DNA) and transduction (RNA) level, the fold differences for RNA were much lower than for DNA. This could be due to several reasons. First, because we did a pooled injection, the injected dose of vector was quite low (4.48×10^12^ vg), with most capsids represented at only 4×10^11^ vg each, so transgene RNA levels were quite low in spinal cord (see RT-qPCR, **Fig. 5a**) which could only be reliably detected in the lumbar region. We couldn’t assess the transduction in the thoracic or cervical regions. Therefore, to get the most accurate read on transduction efficiency, these individual candidates should be injected individually in NHPs at a higher dose and packaging an optimized transgene expression cassette, and careful stereological quantitation of transduced cells should be performed. It could be that while the capsids had remarkably enhanced biodistribution compared to AAV9, the dose was too low to allow effective genome concatamerization and robust transduction. Alternatively, some of the capsids may not mediate all of the intracellular steps required for transduction.

While the NGS reads coming from RNA provided evidence that many of the selected capsids were more functional at transduction of spinal cord compared AAV9, we could not determine which cell types were transduced (anti-HA Frataxin staining of spinal cord did not show any signal; data not shown). Thus, to understand if the capsids could transduce the main targets of gene therapy in the CNS, astrocytes and neurons, we used intrathecal lumbar injection of mice as a surrogate. This experiment also tested whether the NHP-selected capsid transduction would occur in the most common preclinical models of human disease, mice. Using a semi-quantitative analysis of spinal cord transduction, we found that these capsids, similar like AAV9, transduce clinically relevant cell types such as neurons and glia. In most cases, transduction efficiency in mice was similar to AAV9, so these capsids may be used for pre-clinical work in mice.

While not a primary objective of the study, we also assessed AAV encoded DNA barcode frequency for the pooled capsids in brain and peripheral nerves. From these data, certain capsids appear to outperform AAV9 in biodistribution to certain brain regions and nerves. This includes capsid DH1 in brain and DH1, DP1, and DR2 in nerves. It may be of future interest to test these individual capsids in NHPs after intrathecal injection to assess transduction of brain and nerves.

In summary, we have identified capsids with enhanced biodistribution and transduction of spinal cord in NHPs compared to AAV9. Coupled with the lower liver biodistribution and transduction, we consider the four capsids described here, NL1, TH1, TP1, and TR2, all to be preclinical gene delivery candidates to be further validated in NHPs and in mouse models of spinal cord injury and disease.

## Acknowledgements

We thank the research team and especially Alan LaRochelle at Biomere Biomedical Research Models (Worcester, MA) for carrying out the animal studies. We thank Dr. Jessica Chichester of the University of Pennsylvania Human Immunology Core for performing the AAV9 neutralization assays.

We thank the team led by Dr. Encrico Radaelli at the Penn Vet Comparative Pathology Core for performing the core service of IHC staining of spinal cords of mice injected intrathecally with AAV vectors used to generate data for Figure 6.

For generation of the artwork in the figures, BioRender software was utilized for some of the objects. For the AAV capsid cartoons, we used the Protein Data Bank (RCSB PDB) AAV9 capsid structure PDB ID: pdb7MT0^28^. We exported the AAV9 structure in 3D viewer into a drawing program for custom coloring.

## Data Availability Statement

The AAV rep/cap plasmids containing the inserts for the capsids described in the manuscript as well as the electronic sequence maps are available upon completion of a standard Material Transfer Agreement with The Massachusetts General Hospital. Any other raw data and raw sequencing files that support the findings of this study are available from the corresponding author.

## Code availability statement

All code written and used in this study can be freely accessed by emailing the authors, who will provide scripts and methods of use.

## Methods

### Animal experiments

Animal studies were performed at Biomere-Biomedical Research Models (Worcester, MA) according to animal use guidelines and approved procedures. The Test Facility is accredited by the Association for the Assessment and Accreditation of Laboratory Animal Care, International (AAALAC) and registered with the United States Department of Agriculture (USDA) to conduct research in laboratory animals.

#### Non-human primates

A total of four cynomolgus monkeys (*Macaca fascicularis*) were used in this entire study (see Table I for complete animal information).

#### Mouse experiments

A total of 30 adult male C57BL/6 mice (Jackson Laboratories) were utilized for the data described in Figure 6.

### Cell culture

Human 293T cells were obtained from American Type Culture Collection (Manassas, VA). Cells were cultured in high glucose Dulbecco’s modified Eagle’s medium containing HEPES (Invitrogen, Carlsbad, CA) supplemented with 10% fetal bovine serum (FBS) (Sigma, St. Louis, MO) and 100 U/mL penicillin, 100 μg/mL streptomycin (Invitrogen) in a humidified atmosphere supplemented with 5% CO2 at 37 °C.

### AAV plasmid constructs

The iTransduce plasmid, pAAV-CBA-Cre-mut/p41-Cap9 (containing the insert with 7mer peptide sequences has been previously described^16^.

pAAV-CBA-Floxed-STOP-H2B-mPlum plasmid was constructed by digesting the AAV ITR containing plasmid, AAV-CBA-WPRE, after the CBA promoter with HindIII and NheI restriction sites. A DNA fragment for a floxed transcription stop transcription site (3 copies of SV40 late poly A sequence) followed by a H2B protein fused to mPlum was synthesized by Genscript (Piscataway, NJ) and inserted into pUC57-Kanamycin plasmid. Next pUC57-Kan with Floxed-STOP-H2B-mPlum was digested with HindIII and NheI and inserted into the above digested AAV plasmid to create p pAAV-CBA-Floxed-STOP-H2B-mPlum (plasmid electronic map available upon request).

pAAV-CBA-hFrataxin-HA-BC-WPRE plasmid was constructed by digesting the AAV transgene expression plasmid, pAAV-CBA-GFP, with AgeI and NheI to remove the GFP cDNA fragment. A double stranded DNA gBlock™ fragment was ordered from Integrated DNA Technologies (IDT, Coralville, Iowa) which contained human frataxin cDNA fused with a hemagglutinin (HA) tag on its C-terminus. This fragment was inserted into the pAAV plasmid above using the Gibson Assembly® Master Mix (New England Biolabs, Ipswich, MA). This “base” plasmid, pAAV-CBA-hFrataxin-HA was used as the acceptor plasmid to insert barcode fragments downstream of the HA sequence (plasmid map available on request). To insert the unique DNA/RNA barcode sequences into the base plasmid, pAAV-CBA-hFrataxin-HA was digested with XhoI. Next, we ordered double stranded DNA gBlock™ fragments (130 bp) from IDT which had unique 21mer bp barcodes (for 14 capsid candidates as well as AAV9) with flanking homology arms.

These 15 different barcodes were individually cloned into the XhoI-digested pAAV-CBA-hFrataxin-HA plasmid using Gibson Assembly as before. Each barcoded pAAV-CBA-hFrataxin-HA plasmid was complete-plasmid sequenced at PlasmidSaurus (Eugene, OR). AAV9 capsid candidates containing unique 7mer peptides were individually cloned into the AAV9 rep/cap plasmid, pAR9, as previously described^16^.

### AAV library production

Production of the AAV9 iTransduce library was performed as previously described^16^ with some changes. Fifty tissue culture dishes (15 cm diameter) were used (1.5×10^7^ 293T cells seeded per plate), with cells cultured in DMEM containing 10% fetal bovine serum (FBS) and 100 U/mL of penicillin, 100 µg/mLstreptomycin, and 292 µg/mL L-glutamine (Invitrogen). Twenty-four hours after plating, cells were triple transfected using polyethylenimine (PEI) transfection reagent, adding to each 15 cm plate: pAAV-CBA-Cre-mut/p41-Cap9 (containing the insert with 7mer peptide sequences; 200 ng), pAR9-Cap9-stop/AAP/Rep (provides rep in trans, 7 μg), and pAdΔF6 (helper plasmid, 15 μg), and sheared salmon sperm DNA (6 μg). Seventy-two hours post transfection, AAV was isolated from a pool of clarified cell lysate and polyethylene glycol (PEG)-precipitated vector from the conditioned media. The pooled virus was then purified by iodixanol density-gradient ultracentrifugation. Buffer exchange to phosphate buffered saline (PBS) containing 0.001% Pluronic F-68 (Gibco) was done using ZEBA spin columns (7K MWCO; Thermo Fisher Scientific) and further concentration was performed using Amicon Ultra 100kDa MWCO ultrafiltration centrifugal devices (Millipore). We quantified AAV vector genomes (vg) in AAV preparations using TaqMan qPCR with BGH polyA-sequence specific primers and probe^29^. Endotoxin for both vector preparations was determined to be <1 EU/mL using Endosafe® LAL Cartridges and the Endosafe® nexgen PTS™ device (Charles River, Charleston, SC). Vector purity was assessed by silver staining of SDS PAGE gels in which 1×10^10^ vg of each vector was run/lane. Purity of both preparations were >90%. Vector was stored at -80°C until use.

### AAV-CBA-Floxed-STOP-H2B-mPlum production

This vector was produced, purified, and titered by Vector Biolabs (Malvern, PA). Endotoxin level was below 1 EU/mL.

### Barcoded AAV capsid candidate production

We individually packed the AAV-CBA-hFrataxin barcoded genome in each capsid candidate and the AAV9 capsid. For each vector we transfected five tissue culture dishes (15 cm diameter, 1.5×10^7^ 293T cells seeded per plate), with cells cultured in DMEM containing 10% fetal bovine serum (FBS) and 100 U/ml of penicillin, 100 µg/mL streptomycin, and 292 µg/mL L-glutamine (Invitrogen). Twenty-four hours after plating, cells were triple transfected using polyethylenimine (PEI) transfection reagent, adding to each 15 cm plate: pAAV-CBA-hFrataxin-HA-barcode (6 μg), pAR9-peptide of interest (7 μg), and pAdΔF6 (helper plasmid, 15 μg). Seventy-two hours post transfection, AAV was isolated from a pool of clarified cell lysate and polyethylene glycol (PEG)-precipitated vector from the conditioned media. The pooled virus was then purified by iodixanol density-gradient ultracentrifugation. Buffer exchange to phosphate buffered saline (PBS) containing 0.001% Pluronic F68 (Gibco) was done using ZEBA spin columns (7K MWCO; Thermo Fisher Scientific) and further concentration was performed using Amicon Ultra 100kDa MWCO ultrafiltration centrifugal devices (Millipore). We quantified AAV vector genomes (vg) in AAV preparations using TaqMan qPCR with BGH polyA-sequence specific primers and probe^29^. We next pooled the barcoded vectors and endotoxin was determined to be <1 EU/mL using Endosafe® LAL Cartridges and the Endosafe® nexgen PTS™ device (Charles River). Pooled vector purity was assessed by staining SDS PAGE gels with GelCode™ Blue (ThermoFisher) in which 1×10^11^ vgs were run/lane. The purity of both preparations were >90%. Vector was stored at -80°C until use.

### Non-human primate library selection

Animals were considered acclimated to the environment at the time of transfer to the study. Prior to the study, serum from each animal was tested for neutralizing antibodies to AAV9 by the University of Pennsylvania Gene Therapy Program Immunology Core run by Dr. Jessica Chichester. Titers for all animals were at or below 1:5.

#### Round 1 selection

On Day 0, the animal received an IT dose of unselected AAV library as appropriate to group (per the study design **Table I**). The dose volume injected was 0.74 mL and vector dose of 9×10^11^ vg. The animal was given Buprenorphine (0.03 mg/kg, IM) and Meloxicam (0.2 mg/kg, SC) prior to the procedure for the purpose of analgesia. The animal was sedated with Ketamine 7.5-12 mg/kg and Dexdomitor 0.01-0.03 mg/kg mixture, IM; Atipamezole (0.1-0.3 mg/kg, IM) was used for reversal. The animal was positioned in lateral recumbency while on a circulating warm water blanket and/or forced warm air blanket during the procedure. The head was kept in line with the spine and the hips and shoulders were perpendicular to the table. The lower back was arched to increase spacing between the spinous processes. The lumbosacral region (the area over ∼L4/5 for cynomolgus) was clipped and aseptically prepared utilizing 3 alternating scrubs of either povidone iodine or chlorhexidine scrub solution and sponges soaked in 70% isopropyl alcohol. A line block (i.e., Lidocaine/Bupivacaine) ∼0.20-0.50 mL, SC was administered at the lumbar puncture site. A final application of ChloraPrep™ or appropriate antimicrobial was applied to the puncture site and allowed to dry.

The wings of the ileum were palpated to provide anatomical landmarks. The two spinous processes were identified, and in between which the spinal needle (22 g x 1.5") was introduced. The skin was penetrated and needle slowly advanced. After confirmation of placement in the intrathecal space, ∼0.5 mL of CSF was removed prior to dose administration. Next, the AAV vector was slowly administered over 1-2 minutes. After administering the dose, the syringe and needle was left in place for ∼5 seconds and after removal, pressure applied to the injection site. Parameters were observed constantly throughout the procedure including heart rate, respiratory rate and oxygen saturation. Animals were recovered from anesthesia and moved to a recovery area, placed on a circulating warm water blanket and/or forced warm air blanket and covered with a dry towel. Animals were observed following the procedure and kept in the recovery area until the animal was conscious and able to hold itself in a sitting position. Animals were then transported to their home cage. Approximately 2 hours post vector dosing, animals were immunosuppressed with intramuscular dosing of 0.5 mg/kg of dexamethasone, which continued daily until necropsy at day 21.

#### Round 2 selection

The IT injection and animal care was identical to round 1 with the following changes. First the animal was lumbar-injected with the recovered and packaged round 1 library at a dose of 2×10^11^ vg in a volume of 0.6 mL. CSF was collected before injection. Three hours later, the animal was IT-injected in a different lumbar region than the first dose with 3×10^13^ vg of AAV9-CBA-FLOXED-STOP-H2B-mPlum vector in a total volume of 0.6 mL.

#### NHP Necropsy and tissue processing

On Day 21 ±1 day, animals were anesthetized with Ketamine 7.5-12 mg/kg and Dexdomitor 0.01-0.03 mg/kg mixture. Nembutal was administered at 15-30 mg/kg. Once deeply anesthetized, the animal was perfused via left cardiac ventricle with cold heparinized (100 U/mL) saline until the outflow ran clear. Approximately 1 L of heparinized saline was used with a perfusion pump set to ∼400 rpm. Euthanasia was performed per AMVA guidelines. For rounds 1 and 2, flash frozen samples were collected for the entire spinal cord and brain and stored at -80°C.

### DNA isolation from NHP tissue during selection rounds

#### Whole tissue isolation

Spinal cord samples (0.5 cm-1 cm in length) were homogenized using 1.4 mm ceramic beads in a BeadBug tissue homogenizer (Benchmark Scientific, Sayreville, NJ) in buffer ATL of the DNeasy Blood and Tissue Kit (Qiagen). After homogenization, we followed the manufacturer’s instructions to purify DNA. DNA concentration was determined using a NanoDrop spectrophotometer (ThermoFisher).

#### Nuclei isolation and flow sorting

Spinal cord samples were dissociated using a GentleMACS Dissociator (Miltenyi Biotec, Westphalia, Germany) using Nuclei Extraction Buffer (Miltenyi) and homogenate filtered through a 30 µm Smart Strainer (Miltenyi). Nuclei were next purified using a 30% OptiPrep™ iodixanol (Axis-Shield, Oslo, Norway) gradient. The homogenate containing nuclei was layered on top of the iodixanol gradient and centrifuged at 10,000x*g* at 4°C. The gradient was carefully aspirated leaving the nuclei pellet in the tube. Nuclei were resuspended in 500 µl of cold PBS. Next nuclei were labeled with Vybrant™ DyeCycle™ Violet Stain (ThermoFisher) and were sorted on a BD FACS Aria II Cell Sorter (Becton Dickinson, Franklin Lake, NJ). First, violet stain positive nuclei were gated on (PI) using Pacific Blue-A laser. Then singlets were selected using FSC-A and FSC-H axes (P2). Finally, we selected mPlum-positive nuclei using a PE-Cy5-A laser. Nuclei (mPlum-positive or negative) were sorted into tubes and stored at -80°C until DNA isolation. For DNA isolation, we used the Arcturus® PicoPure® DNA Extraction Kit (ThermoFisher).

### Next generation sequencing of library

Next generation sequencing was performed on the plasmid AAV9 library pool, as well as following packaging of capsids. Sequencing was also performed following PCR rescue of the cap gene fragment (either from NHP spinal cord tissue or from nuclei sorted by flow cytometry). For each round of selection vector DNA corresponding to the insert-containing region was amplified by PCR using either Phusion High-Fidelity enzyme or Q5 polymerase (both from New England Biolabs using Forward primer: 5’-AATCCTGGACCTGCTATGGC-3’, and reverse primer: 5’-TGCCAAACCATACCCGGAAG-3’). PCR products were purified using a QIAquick PCR Purification Kit (Qiagen). Unique barcode adapters were annealed to each sample, and samples were sequenced on an Illumina MiSeq (150bp reads) at the Massachusetts General Hospital Center for Computational and Integrative Biology DNA Core. Approximately 50,000-100,000 reads per sample were analyzed. Sequence output files were quality-checked initially using FastQC (http://www.bioinformatics.babraham.ac.uk/projects/fastqc/) and analyzed on a program custom-written in Python. Briefly, sequences were binned based on the presence or absence of insert; insert-containing sequences were then compared to a baseline reference sequence and error-free reads were tabulated based on incidences of each detected unique insert. Inserts were translated and normalized.

### Non-human primate candidate barcoded library screen

Two adult male cynomolgus monkeys (see Table I for NHP information) were intrathecally injected in the same manner as for the selection, except with 4.48×10^12^ vg of the pooled barcoded capsids. Three weeks later, animals were perfused with sterile heparinized saline. Samples for paraffin embedding were fixed in 10% neutral buffered formalin before transferring the samples to PBS. Samples for cryosectioning were fixed in 4% buffered formaldehyde for 24 h before transferring samples to PBS. Samples for DNA and RNA extraction were immediately frozen on dry ice and stored at -80°C.

### DNA and RNA isolation from NHP whole tissue for barcoded AAV capsid candidates

#### DNA isolation

Spinal cord (and other tissue) samples (0.5 cm-1 cm in length) were homogenized using 1.4 mm ceramic beads in a BeadBug tissue homogenizer (Benchmark Scientific) in buffer ATL of the DNeasy Blood and Tissue Kit (Qiagen). After homogenization, we followed the manufacturer’s instructions to purify DNA. DNA concentration was determined using a NanoDrop spectrophotometer (ThermoFisher Scientific).

#### RNA isolation

Spinal cord (and other tissue) was placed into Qiazol (Qiagen) and subjected to homogenization with 1.4 mm ceramic beads in a BeadBug tissue homogenizer (Benchmark Scientific). After performing a phenol/chloroform extraction of RNA, the RNA was precipitated using isopropranol and centrifugation, followed by a 75% ethanol wash. The RNA pellet was resuspended in RNase-free water and stored at -80C until further processing. The RNA sample was further purified using RNeasy spin columns (Qiagen) and the sample was eluted in RNase-free water. Next residual DNA contamination was removed using DNaseI treatment with a DNA-*free™* kit (Ambion by Life Technologies). cDNA was synthesized from RNA (approximate input 500 ng of RNA) using the SuperScript™ IV VILO™ Master Mix reverse transcriptase (RT) kit. For each sample we did a control where the RT enzyme was left out to ensure the PCR amplicons originated from RNA and not contaminated DNA in the RNA samples.

### Quantitative PCR to measure absolute amounts of AAV genomes and transgene mRNA in spinal cord of pooled AAV injection in NHPs

DNA and RNA isolation and cDNA synthesis from NHP spinal cord was performed as described immediately above. To determine the amounts of AAV vector genomes or transgene expression (frataxin-HA) we used the same Taqman probes and primers targeting BGH polyA used to titer our AAV vectors. For AAV vector genomes, we used a standard curve with an AAV plasmid to perform absolute quantitation. We used 54 ng of genomic DNA template from lumbar, thoracic, and cervical spinal cord as input for the qPCR. A separate qPCR reaction for each sample was performed to detect the NHP gene UBE2D2 (ubiquitin conjugating enzyme E2 D2) using the Taqman Gene Expression Assays 20x mix (Catalog #4351372, assay ID Mf07285893_s1, ThermoFisher Scientific). AAV genomes were inferred from the AAV plasmid standard curve and adjusted to AAV genomes/µg of genomic DNA. For each AAV genome sample we calculated the 2^ΔCt^ (sample Ct – sample with lowest Ct) of the UBE2D2 qPCR sample and then normalized the AAV genomes to that value. This compensated for DNA input differences for each sample.

For the RT-qPCR samples, we used the AAV plasmid as a standard curve as above to calculate absolute amounts of cDNA. We used 1 µl of the RT reaction from mRNA isolated from NHP lumbar spinal cord as template for the qPCR. A separate qPCR reaction for each sample was performed to detect the NHP cDNA of GAPDH (glyceraldehyde-3-phosphate dehydrogenase) using the Taqman Gene Expression Assays 20x mix (assay ID <u>Mf04392546_g1</u>, ThermoFisher Scientific). This probe spans exons, so it is selective for cDNA detection over genomic DNA. cDNA values for AAV transgene were normalized to the respective GAPDH Ct values for each sample to control for differences in input total cDNA. All qPCR reactions were performed in a 7500 FAST qPCR system (Applied Biosystems).

### PCR to amplify barcodes for NGS

For both the DNA and RNA samples from the pooled barcoded library injected animals we used the following reagents. A PCR using either the DNeasy-purified DNA or the cDNA from the RT reaction as template was performed (usually 150 ng DNA template input). We used Q5 polymerase (NEB) and primers that amplify a 182bp amplicon which contains the unique barcode for each capsid. Primers were: SC-Lib-Fwd (5’ GATGCTTACCCTTACGACGT 3’) and SC-Lib-Rev (5’ CAGCGTATCCACATAGCGTA 3’). The PCR product was purified using the PureLink™ PCR Purification Kit (ThermoFisher Scientific) and samples were submitted for NGS as described above. We initially sequenced an aliquot of the pooled barcoded library to ascertain the precise frequencies of the barcoded variants on input. Output frequencies in all tissues were normalized to these initial ratios. Insert-containing sequence reads were binned using Hamming distance, comparing to the known barcode sequences; additional quality control was undertaken to ensure only true barcoded reads were assigned to each capsid.

### Intrathecal lumbar injection in mice

Adult (6-8 weeks old) male C57BL/6 mice received a single intrathecal dose of each of the four vector groups (n=5/group) described in Figure 6 or control treatment (vector formulation buffer). Animals received a prophylactic dose of Buprenorphine SR (1 mg/kg, SC), then were anesthetized by isoflurane to effect. Syringes were loaded individually prior to each dose with 10 μL of dosing solution per animal. The test material was delivered manually, using an insulin syringe with a 28-30-gauge needle, by intrathecal injection to the lumbar spine at the L4-L5 or L5-L6 intravertebral space. The tip of the needle was introduced into the lumbar spine, and proper insertion of the needle was confirmed by tail flick reflex. After slowly injecting the total dose volume, the needle will be allowed to stay in place for a minimum of 5 seconds to avoid efflux of the test material to periphery. Animals were placed on a circulating water heating pad until fully recovered from anesthesia. Four weeks after administration, animals were euthanized (carbon dioxide asphyxiation, to effect, in conjunction with exsanguination), and tissues were removed for fixation (histology) or snap frozen in liquid nitrogen.

### Immunohistochemical staining of mouse tissues from mice injected with AAV capsids packaging AAV-CAG-GFP cassette

Tissues designated for histologic processing were placed in appropriately sized histology cassettes and fixed in 10% NBF (neutral buffered formalin) at ambient temperature for 24-48 hours then transferred to 1X PBS and stored at 5 °C ± 3 °C until embedded.

The immunohistochemical staining of GFP in spinal cords and livers of the injected mice was performed by the Penn Vet Comparative Pathology Core (University of Pennsylvania, Philadelphia, PA). Briefly, the formalized fixed tissue as paraffin embedded and 5 µm thick paraffin sections mounted on ProbeOn™ slides (Thermo Fisher Scientific). The immunostaining procedure was performed using a Leica BOND RX^m^ automated platform combined with the Bond Polymer Refine Detection kit (Leica DS9800). Sections on slides were dewaxed and rehydrated in deionized water and then pretreated with the epitope retrieval BOND ER2 high pH buffer (EDTA Based pH=9.0, Leica AR9640). Next endogenous peroxidase was inactivated with 3% H2O2. Nonspecific protein-protein interactions were blocked with Leica PowerVision IHC/ISH Super Blocking solution (Leica PV6122) at RT. The primary rabbit polyclonal anti-GFP antibody (Cat. no. A11122, ThermoFisher) was diluted 1/1000 using Cell Signaling Technology (CST) diluent solution and was applied for 45 minutes at RT. A biotin-free polymeric IHC detection system (Leica DS9800) consisting of HRP conjugated Gt anti-Rb IgG secondary antibody was applied for 25 minutes at RT. Immunoreactivity was detected with the diaminobenzidine (DAB) chromogen reaction. Slides were counterstained in hematoxylin, dehydrated in ethanol series, cleared in xylene, and mounted with resinous mounting medium (Thermo Scientific, ClearVue™ coverslipper). Examination and grading of the GFP staining in cell types (e.g. neurons, glia) was performed by trained personnel at the core facility.

### Data analysis

To assess common amino acids across the 7-mer insert between the 13-chosen candidates, we used the web-based application WebLogo (https://weblogo.threeplusone.com/)^30, 31^ which generates alignments of the input sequences, whereby the height of each amino acid corresponding to its frequency at that position.

### Statistics

We used GraphPad Prism 9.0 for PC for statistical analysis. For comparison of biodistribution (AAV genomes) between animals and across regions we used an ANOVA followed by a Šídák’s multiple comparisons test. To compare cDNA levels of AAV transcripts between NHPs #1001 and #1002 in the lumbar region, we used an unpaired two tailed t-test; p values <0.05 were accepted as significant.

