## Supplementary Figures for "*In vivo* selection in non-human primates identifies superior AAV capsids for on-target CSF delivery to spinal cord"

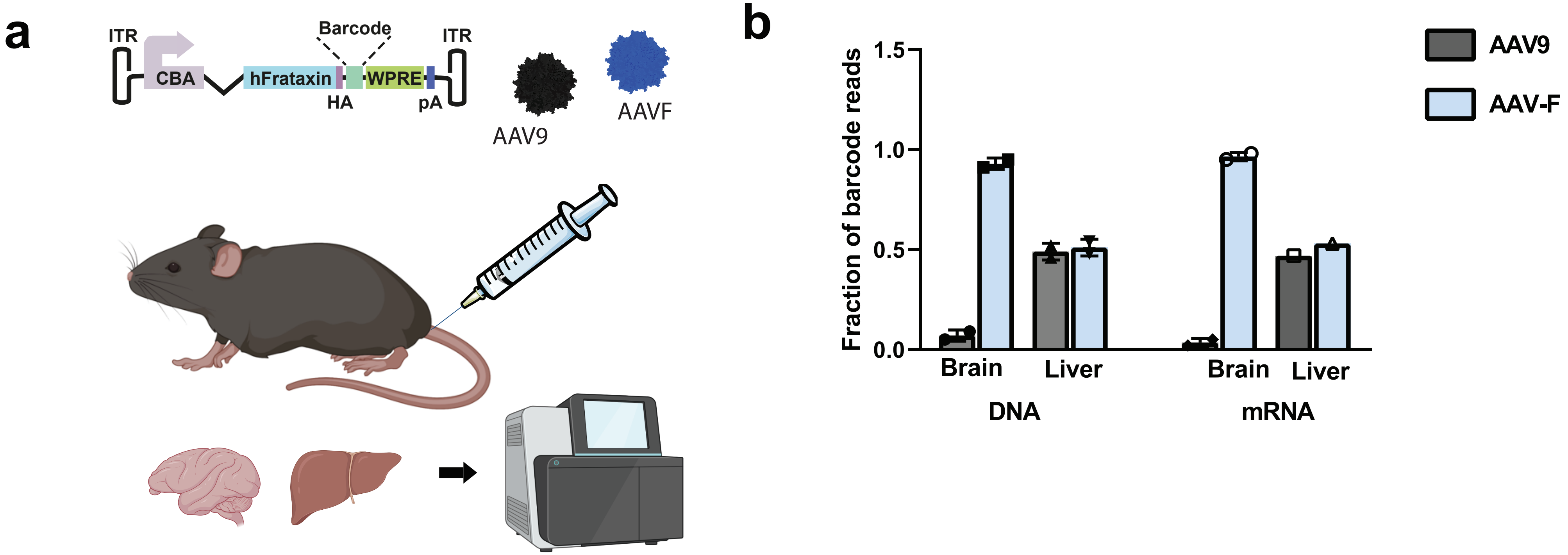


**Fig. S1. Validation of barcoded AAV genomes to detect differences in biodistribution and transgene expression of validate capsids.** An AAV expression plasmid was constructed with a CBA promoter, hFrataxin cDNA with a HA tag, and a DNA barcode that can be measured at the DNA and RNA levels. Unique barcodes were made for AAV9 and AAV-F and packaged into these capsids.
Mice (n=2) received a 1:1 mixture of the capsids by tail vein injection, and liver and brain were harvested for measurement of barcode reads by NGS for both DNA and RNA.Quantitation of reads obtained in the liver and brain. Biological replicates are shown for each group.
Only one mouse is shown for liver mRNA reads due to a contamination issue with one of the mouse livers. All other samples show two biological replicates. Error bars depict the standard deviation from the mean.

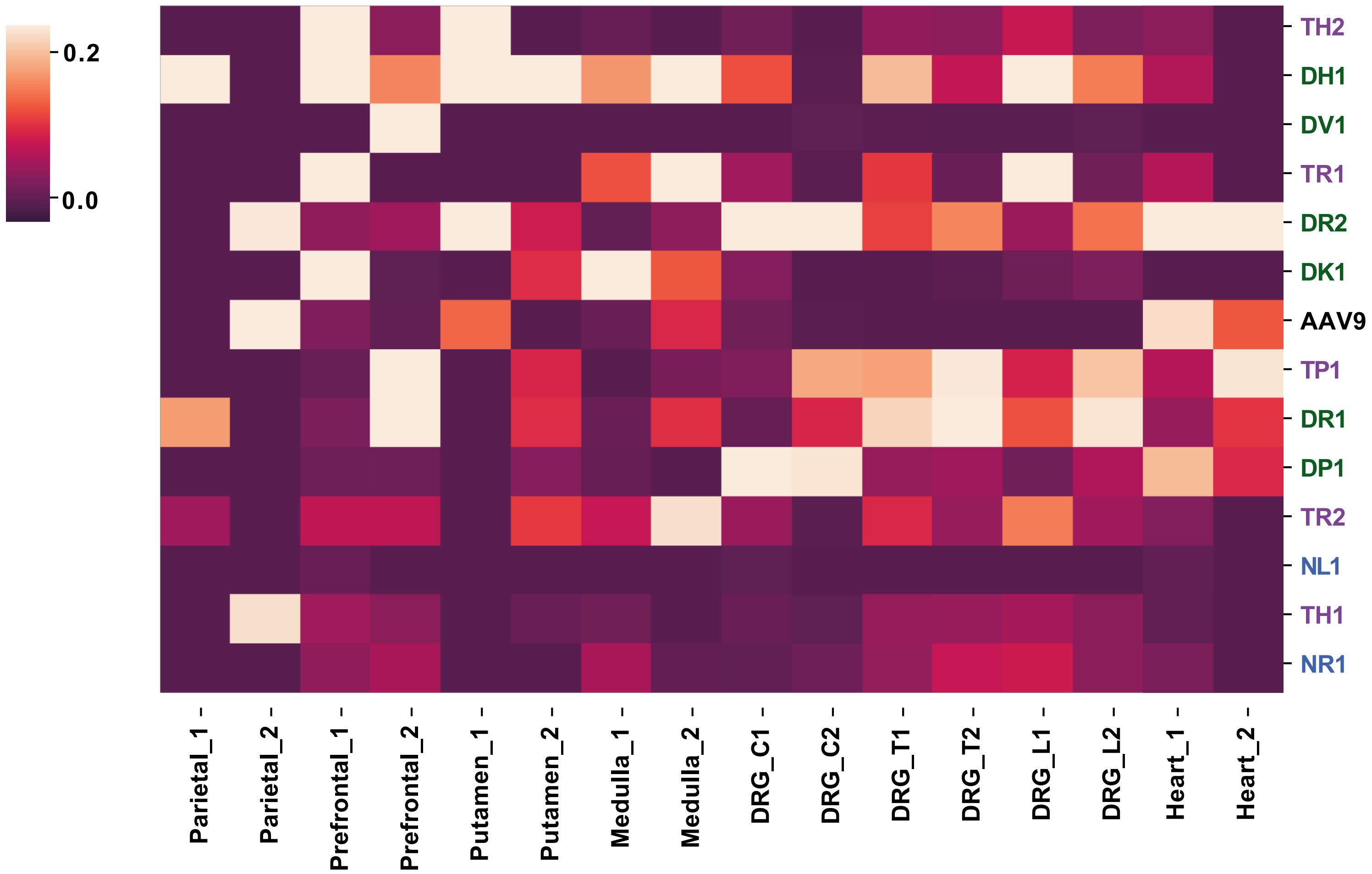


**Figure S2: Variant frequency in non-spinal cord tissues of NHPs injected intrathecally with barcoded AAV capsid candidates.** Heatmap showing relative frequency of each AAV variant in various regions of the CNS, along with heart and liver. Variants are clustered by expression pattern. Number (1, 2) refers to each animal. Parietal, parietal lobe. Prefrontal, prefrontal cortex. DRG, dorsal root ganglion.

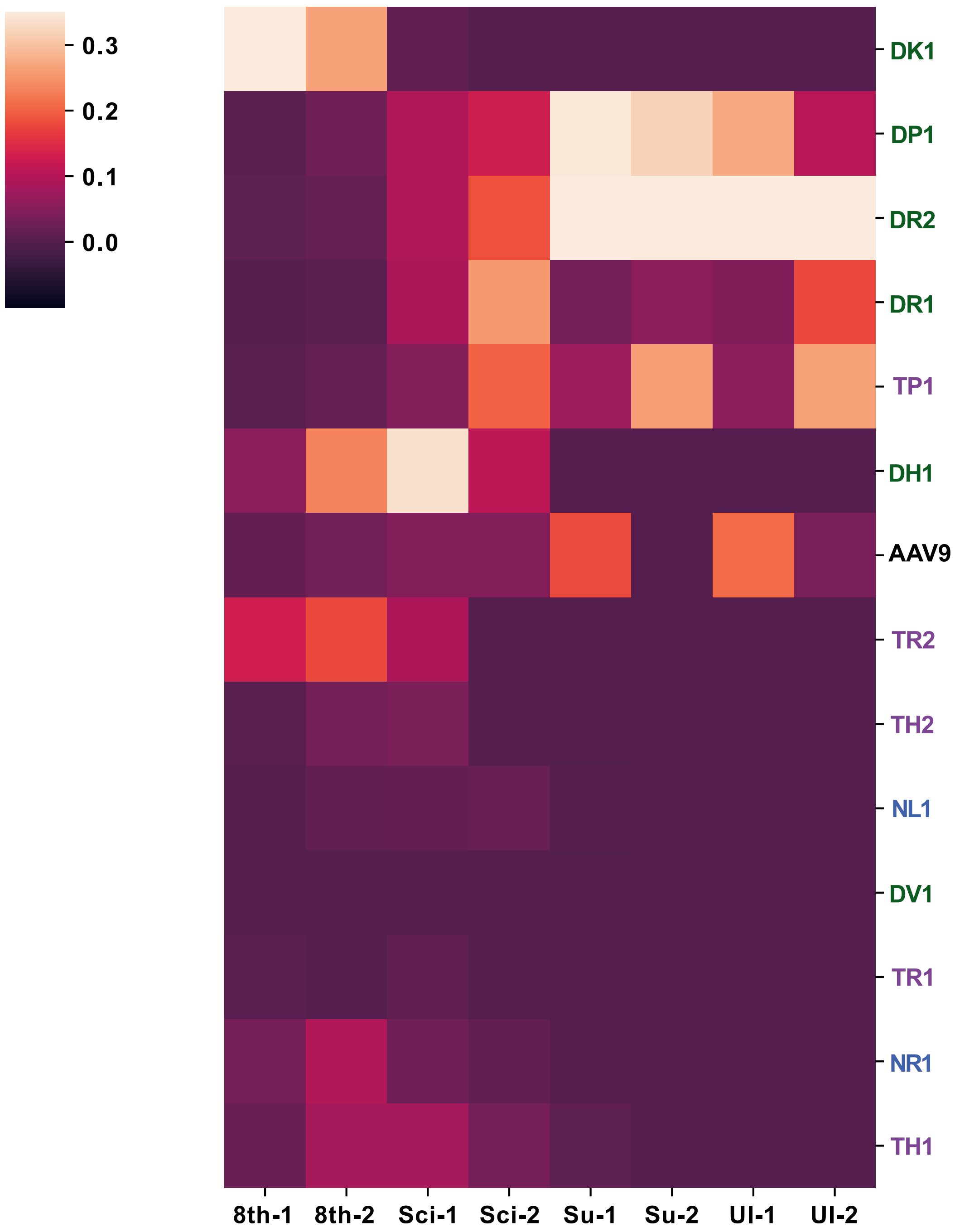


**Figure S3: Variant frequency in peripheral nerve tissues of NHPs injected intrathecally with barcoded AAV capsid candidates**. Heatmap showing relative frequency of each AAV variant in peripheral nerves. Variants are clustered by expression pattern. Number (1, 2) refers to each NHP (#1001 and #1002). 8th, 8th cranial nerve. Sci, sciatic nerve. Su, sural nerve. Ul, ulnar nerve.

**a**

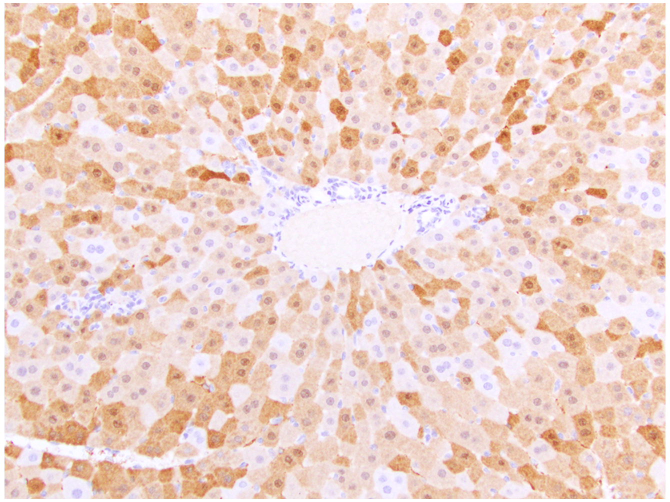

**b**

| Grade | % positive cells |
| --- | --- |
| 5 | 76-100% |
| 4 | 51-75% |
| 3 | 26-50% |
| 2 | 1-25% |
| 1 | <1% |
| 0 | 0% |

**c**

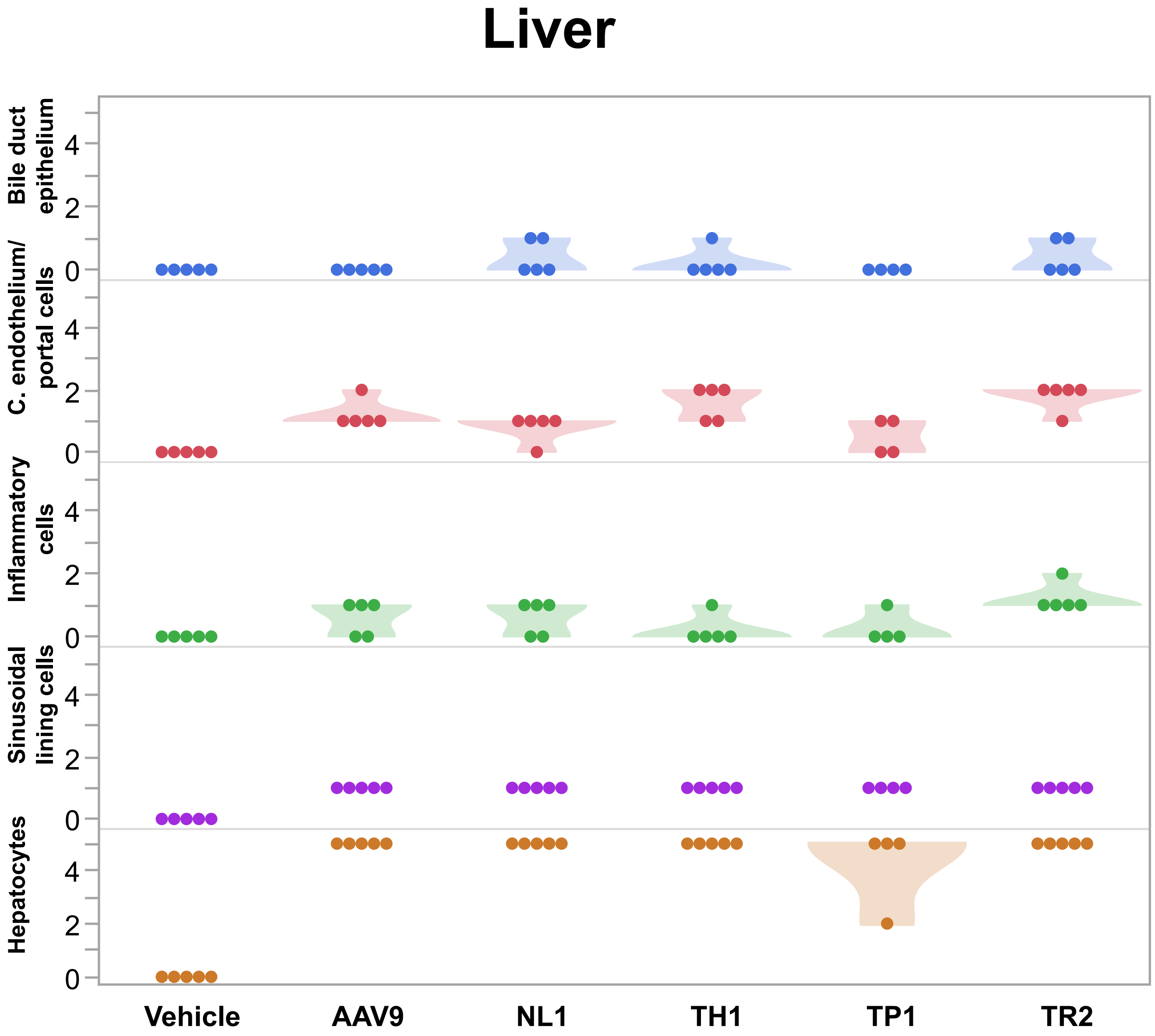


**Figure S4. Transduction of mouse liver by NHP selected AAV capsid candidates after lumbar intrathecal bolus injection.** Mice (n=5/ capsid) were injected intrathecally with AAV9, NL1, TH1, TP1, and TR2. All capsids were injected at $8.3x10^{10}$ vg/an with the exception of NL1, which was dosed at 4.8 $x10^{10}$ vg/an **a.** Sample image of transduced liver. Brown DAB (3,3'-Diaminobenzidine) staining indicates GFP immunoreactivity. **b.** scoring grading for transduction efficiency. **c.** Transduction scoring for various cell types in the mouse liver. Each data point represents one mouse.
